# Which way is the bookstore? A closer look at the judgments of relative directions task

**DOI:** 10.1101/391326

**Authors:** Derek J. Huffman, Arne D. Ekstrom

**Affiliations:** Center for Neuroscience, University of California, Davis; Department of Psychology, University of California, Davis; Neuroscience Graduate Group, University of California, Davis; Psychology Department, University of Arizona

**Keywords:** cognitive map, judgments of relative direction (JRD), map drawing task, map sketching, group-level nonparametric permutation test

## Abstract

We present a detailed analysis of a widely used assay in human spatial cognition, the judgments of relative direction (JRD) task. We conducted three experiments involving virtual navigation interspersed with the JRD task, and included confidence judgments and map drawing as additional metrics. We also present a technique for assessing the similarity of the cognitive representations underlying performance on the JRD and map drawing tasks. Our results support the construct validity of the JRD task and its connection to allocentric representation. Additionally, we found that chance performance on the JRD task depends on the distribution of the angles of participants’ responses, rather than being constant and 90 degrees. Accordingly, we present a method for better determining chance performance.

## 1 Introduction

Navigation is critical to the survival of many organisms. The ability to remember the locations of goal-relevant landmarks—e.g., the location of a blueberry bush relative to one’s dwelling—allows the navigator to adapt to its environment and behave in a goal-oriented manner. The locations of such goal-relevant landmarks are inherently relational, and thus these spatial relationships are typically described in terms of relative directions and distances. In fact, the past several decades have seen the advancement of several behavioral techniques for studying human spatial memory in the laboratory, and these techniques typically assay participants’ knowledge of the relative directions or relative distances (or both) between landmarks. Some of the most commonly used tasks fit broadly into the following categories: 1) pointing tasks (which provide a measure of relative directions; e.g., Waller & Hodgson, 2006), 2) distance estimation tasks^1^ (which provide a measure of relative distances; e.g., Richardson, 1981), 3) map drawing tasks (which provide a measure of both relative directions and distances; MacKay, 1976).

Pointing tasks are a valuable way to assess spatial knowledge without the confound of the participant continuously changing position while doing so. Another advantage is that pointing tasks are readily adapted to computerized testing, and thus can be employed during neuroimaging. One example of a pointing task involves participants pointing to the direction of landmarks while they are oriented in an environment—i.e., these tasks ask participants to point to landmarks relative to their current egocentric position and heading. These tasks have been referred to as the “egocentric pointing task” (Waller & Hodgson, 2006) and the “scene and orientation-dependent pointing (SOP) task” (Zhang, Copara, & Ekstrom, 2012; Zhang, Zherdeva, & Ekstrom, 2014; Ekstrom, Arnold, & Iaria, 2014; Ekstrom, Huffman, & Starrett, 2017)^2^. There is also a pointing task, dubbed “the judgments of relative direction (JRD) task” (Rieser, 1989; Shelton & McNamara, 2001; Waller & Hodgson, 2006; Zhang et al., 2012, 2014; Ekstrom et al., 2014, 2017), in which participants are asked to recall the locations and directions of landmarks relative to each other, irrespective of their current egocentric position and heading—i.e., pointing to landmarks relative to imagined positions and headings. There are several versions of the JRD task, but the most typically used version asks participants to imagine themselves standing near a landmark, facing another landmark, and to point in the direction of a third landmark—i.e., “Imagine you’re standing at A, facing B. Please point to C.” (Shelton & McNamara, 2001; Waller & Hodgson, 2006; Zhang et al., 2012, 2014; Ekstrom et al., 2014, 2017).

The SOP task is thought to tap into cognitive representations of locations relative to the navigator (Wang & Spelke, 2000, 2002; Waller & Hodgson, 2006), while the JRD task is thought to rely to a greater degree on the flexible use of memory of the locations of landmarks relative to each other (Waller & Hodgson, 2006). Supporting the notion that these tasks rely on partially distinct cognitive processes, Waller and Hodgson (2006) provided evidence of a double dissociation between performance on the SOP task and the JRD task. Specifically, participants learned the location of objects within a room-sized environment prior to entering a chamber at the center of the room. Participants then performed the SOP task and the JRD task under conditions with and without disorientation. Their results indicated that participants performed worse on the SOP task following disorientation, supporting previous findings (Wang & Spelke, 2000, 2002). Interestingly, however, the converse was true for the JRD task—that is, participants actually performed better on the JRD task following disorientation. The authors suggested that these results provide evidence that disorientation causes participants to switch from a high precision, online system to a lower precision memory system involving the relations of positions of objects to each other. Moreover, the finding that performance was better on the JRD task following disorientation suggests that participants were correctly updating their position and facing angle based on their imagined heading rather than based on their actual position in space. Altogether, the results from Waller and Hodgson (2006) demonstrate that the JRD task allows the investigation of the representation of spatial information in memory, and the observed improvement in performance following disorientation suggests that the JRD task favors allocentric forms of representation.

The JRD task, however, particularly when participants need to imagine a new heading that actively conflicts with their current heading (i.e., imagine facing 180 degrees in the room you are currently in without turning), clearly involves some egocentric components (Hintzman, O’Dell, & Arndt, 1981; Rieser, 1989; May, 2004). Somewhat in contrast, map-drawing does not necessitate reproducing first-person facing angles from navigation, and instead, relies largely on one’s memory for the position and inter-relationship of landmarks to each other. In this way, although map drawing does have some memory-related biases (for example, north is up), it is often taken as a comparatively pure measure of an allocentric representation. Participants are often asked to perform map drawing using paper and pencil (e.g., Shelton & McNamara, 2001, 2004a, 2004b; Labate, Pazzaglia, & Hegarty, 2014) and these techniques have more recently been extended to computer-based tasks (e.g., Weisberg, Schinazi, Newcombe, Shipley, & Epstein, 2014). Despite the connection between allocentric reference
 frames, map drawing, and the JRD task, to our knowledge, there have been no direct attempts in the literature to determine how much common variance is shared by map drawing and the JRD task. Moreover, studies that have investigated the relationship between performance on the JRD task and the map drawing task have typically used methods that can only provide indirect evidence of such a relationship.

For example, previous studies have investigated whether the preferred orientation of JRD performance is related to the preferred orientation of participants’ maps—i.e., whether the imagined heading from which JRD performance was the best was also drawn as north on the map drawing task (Shelton & McNamara, 2001, 2004a, 2004b; Shelton & Marchette, 2010; Marchette & Shelton, 2010). Shelton and McNamara (2001, 2004a) found that participants pointed more accurately on the JRD task when the imagined heading was aligned with the originally encoded orientation compared to when the imagined heading was misaligned to this orientation. Additionally, participants tended to draw their map from the originally encoded orientation. These findings generally support the notion that initial egocentric alignment affects performance on both tasks and thus provides indirect evidence that these tasks rely on at least partially overlapping cognitive representations. However, because such effects of imagined facing angle can also be considered a form of encoding specificity (Tulving & Thomson, 1973), in other words, that memory is typically better when retrieved from the same conditions as it was encoded, it is not clear that such findings in fact show commonality in the underlying spatial representations participants employ to perform the two tasks. Additionally, the results of other studies have failed to find a correspondence between the preferred orientation across these two tasks (e.g., Marchette & Shelton, 2010). Furthermore, these analyses address a very specific type of correspondence between the two tasks, leaving open the possibility that performance is related between these tasks but in a manner that cannot be detected in these analyses.

In partial support of the notion that map drawing and JRD performance are related in a way that is not necessarily related to preferred orientation, recent studies have investigated whether there is a relationship between performance on map drawing tasks and both the SOP task (Hegarty, Montello, Richardson, Ishikawa, & Lovelace, 2006; Labate et al., 2014; Schinazi, Nardi, Newcombe, Shipley, & Epstein, 2013; Weisberg et al., 2014) and the JRD task (Bryant, 1984; Schinazi et al., 2013). The results of these studies have generally shown a relationship between how well participants perform on pointing tasks and map drawing tasks (Bryant, 1984; Hegarty et al., 2006; Labate et al., 2014; Weisberg et al., 2014; but see: Schinazi et al., 2013). Additionally, performance on the JRD task (Kozlowski & Bryant, 1977; Bryant, 1982) and the map drawing task (Hegarty et al., 2006; Ishikawa & Montello, 2006; Labate et al., 2014; Weisberg et al., 2014) have been shown to correlate with self-reports of navigational ability, such as sense of direction. Moreover, Waller and Haun (2003) developed a scaling method for reconstructing maps from pointing on a SOP task. They reported good correspondence between such maps and participant-drawn maps—as assessed both by observers’ ratings of these different maps based on viewing them and by the fit of a bidimensional regression analysis—thus supporting the notion that the SOP task contained at least some information employed when drawing maps. Taken together, however, the evidence for correspondence between pointing tasks and map drawing tasks is equivocal (e.g., Marchette & Shelton, 2010; Schinazi et al., 2013), and the evidence provided by these analyses is indirect. Hence, important insight into the nature of the representations employed across these tasks could be afforded by a more direct analysis.

As evidenced above, the JRD task has provided fundamental insight into human spatial cognition. However, several of the underlying properties of the JRD task remain poorly understood. The concept of “construct validity” suggests that a metric should correspond with other measures that we think tap into the same underlying process. The first important question of construct validity we address is whether repetition improves accuracy and confidence in the JRD task. This tests an important assumption that as we emphasize the same spatial information, representations of this information should increase in fidelity. Second, if the JRD task provides insight into consciously accessible knowledge of spatial relationships in an environment, then participants should also show awareness of this fact, and thus we predict that accuracy in the JRD task and confidence should correlate. Third, we assessed the reliability of participants’ responses on the JRD task—i.e., test-retest reliability, an important feature of construct validity (specifically, that responses are not otherwise random or unreliable). Fourth, we tested the hypothesis that judgments of relative direction and map drawing recruit similar cognitive processes involving a form of allocentric representation. We tested this hypotheses using two analytical techniques, namely, a group-level analysis and a within-subjects analysis of the pattern of errors on both tasks. A clearer understanding of relationships in performance, or a lack thereof, between the JRD task and the map drawing task would support the idea that the JRD task measures, to some extent, allocentric knowledge obtained from navigating the environment. Fifth, while previous studies have typically assumed that chance performance on the JRD task was 90 degrees, we tested how we might detect chance performance in the JRD task; unexpectedly, we discovered in the course of this research that chance performance depends on the underlying distribution of participants’ responses on the task. Thus, chance performance is unknown and can vary across participants (and potentially across groups of participants; i.e., different populations). Although, we note that this may depend, to some extent, on details of the experimental design of the JRD questions themselves. Here we present two nonparametric permutation tests for assessing chance performance in a response-independent manner, and our approach can be applied both at the level of individual participants and at the group level.

## 2 Experiment 1

As outlined in the Introduction, there are several questions regarding the nature of the JRD task, and we assessed the first two of those questions in Experiment 1. Specifically, first, we investigated how performance on the JRD task changes over repeated rounds of navigation and testing. Second, we assessed participants’ subjective confidence of their performance, and we tested the hypothesis that individual differences in pointing performance would correlate with differences in subjective performance, which would support the hypothesis that participants are aware of their spatial memory ability.

### 2.1 Method

#### 2.1.1 Participants

Twenty-four participants were recruited from the UC Davis community. Participants consented to the procedures in accordance with the Institutional Review Board of UC Davis and received course credit for their participation. Participants were between 18 and 22 years old (mean = 19.9; 19 females, 5 males). Eight participants were excluded because they failed to perform significantly better than chance (one-tailed *p* > 0.05), as assessed by a permutation test (see 2.1.6 Data Analysis); therefore, 16 participants were included in the main analyses.

#### 2.1.2 Virtual Environment

The experiment was programmed in the Unity game engine (https://unity3d.com). The virtual environment in both experiments consisted of a square arena that spanned 200 × 200 virtual meters (vm). The perimeter of the environment was bounded by brick walls, which were 6 vm tall. The environment contained 10 target stores, and the coordinates of the stores were selected such that the distribution of the imagined headings relative to the main axes of the environment was approximately uniform (calculated with custom-written R code; see the left panel Figure 5 for a map of the locations of the store coordinates [i.e., labeled “Answers”]). For each participant, the 10 stores were randomly drawn from a list of 23 stores—in alphabetical order: 1st Bank, Bakery, Barber, Bike Shop, Book Store, Butcher Shop, Camera Store, Cell Phones Store, Chinese Food Restaurant, Clothing Store, Coffee Shop, Craft Shop, Dentist, Fast Food Restaurant, Florist, Grocery Store, Gym, Ice Cream Shop, Music Store, Pet Store, Pharmacy, Pizzeria, and Toy Store—and each store was randomly assigned to one of the 10 store coordinates. The random assignment of stores controls for semantic effects of the stores across participants (e.g., the Fast Food Restaurant and the Pizzeria are more semantically similar to each other than they are to the Music Store). Each store was the same shape and size (10.9 vm long × 15 vm wide × 6 vm tall) but differed in the colors of the walls and awnings (e.g., brick walls with green awnings) as well as unique objects in the windows (e.g., guitars for the Music Store). The eye height of the avatar was 1.56 vm above the ground.

#### 2.1.3 Storefront Familiarization Task

Prior to beginning the navigation task, participants performed the storefront familiarization task. In this task, they viewed each of the 10 stores for 3 seconds each. Participants were instructed to read its name and remember what the store looked like.

#### 2.1.4 Navigation Task

Participants performed 6 rounds of the navigation task (Figure 1; Caplan et al., 2003; Newman et al., 2007), each of which was followed by a round of the JRD task. During the navigation task, participants were instructed to find each of the 10 stores, the order of which was randomized within each block. Participants indicated that they found a store by running into it, at which point the instructions for the next store appeared. After participants found all 10 stores, the JRD task began.

**Figure 1:**
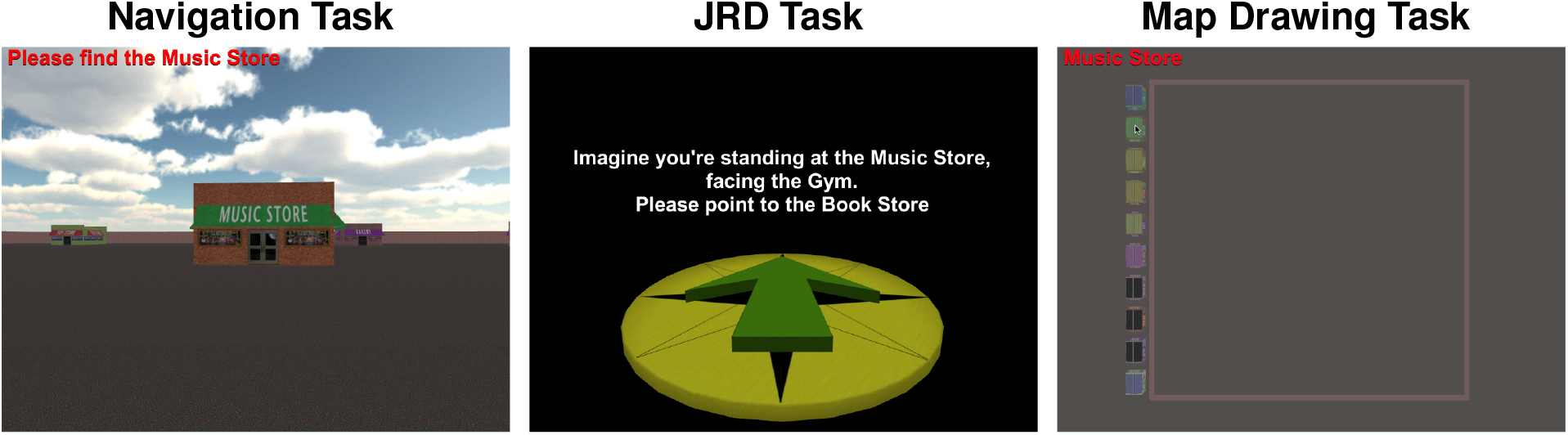
Behavioral tasks. In the navigation task (Left Panel), participants were asked to find each of 10 stores in a random order. In the JRD task (Middle Panel), participants were asked questions of the following structure: “Imagine that you’re standing at Store A, facing Store B. Please point to Store C. “ After selecting their response, participants were asked to indicate their confidence, on a scale of 1 to 4: 1=“very unsure”, 2=“somewhat unsure”, 3=“somewhat sure”, 4=“very sure”. In Experiment 2, participants also performed a map drawing task (Right Panel).

#### 2.1.5 JRD Task

During the Judgments of Relative Direction (JRD) task, participants were asked to imagine that they were standing at one store, facing a second store, and then to point in the direction of a third store—i.e., “Imagine that you’re standing at Store A, facing Store B. Please point to Store C.” Participants used a pointer to indicate the direction of the target store (Figure 1). Questions were presented on a black background to control for sensorimotor alignment effects (i.e., better performance when the imagined heading is aligned with the participant’s actual heading; Kelly, Avraamides, & Loomis, 2007; see also: Hintzman et al., 1981; Rieser, 1989; May, 2004). Questions were randomly sampled from a pool of 10 stores, which gives a total of 720 possible questions (*n* × (*n* − 1) × (*n* − 2), where n is the number of target stores). After selecting their pointing response, they were asked to indicate their confidence, on a scale of 1 to 4: 1=“very unsure”, 2=“somewhat unsure”, 3=“somewhat sure”, 4=“very sure”. Each block consisted of 20 questions.

#### 2.1.6 Data Analysis

Data were analyzed in R (version 3.3.1; R Core Team, 2016) using custom-written code, base R functions, and *ggplot2*. For several of our analyses, we used linear regression models in which we treated participants as a random factor (i.e., random intercepts and slopes), using the *lme4* package (1.1-14; Bates, Mächler, Bolker, & Walker, 2015) and we assessed significance via likelihood ratio tests, using the stats package *anova*.

We used a permutation test to determine whether each participant performed better than chance. This procedure was motived by the results of our simulations (see Simulation of Chance Performance on the JRD Task). For each permutation, the participant’s responses were randomly shuffled and the median absolute angular error was calculated with these arbitrarily associated correct angles. This procedure was repeated 10,000 times per participant. One-tailed nonparametric p values were calculated using the following equation (Ernst, 2004):

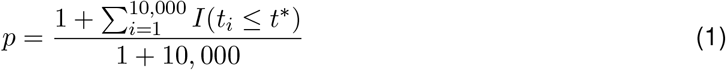

where *I*(·) is an indicator function that takes the value of 1 if true and the value of 0 if false, *t*, is the median error for the *i*th random shuffle, and *t*\* is the observed median error (i.e., the unshuffled value). Note, *t* is used to denote the general test statistic, which in our case was the median angular error. Thus, this equation calculates the proportion of permuted values that lie at least as extreme as the observed value (in the negative direction because we are interested here in testing whether the angular error is significantly less than chance; note, this procedure determines whether the empirical accuracy is in the lower tail of the resultant null distribution). To maximize the similarity between a full permutation and the Monte Carlo permutation test test, 1 is added to both the numerator and the denominator (i.e., in the full permutation, the exact labeling would occur once). We typically advocate for two-tailed nonparametric tests (Huffman & Stark, 2014, 2017); however, in the case of participant exclusion, we propose that it is better to err on the side of conservative exclusion, thus motivating the use of one-tailed tests. However, we have found that participants typically either perform well-above chance (e.g., *p* < 0.001) or at chance with p values well above 0.05, thus the choice of one-or two-tailed tests for participant exclusion might make little difference in practice. Note, here we included all trials in our analysis under the assumption that including as many trials as possible would help to increase the power of detecting significant performance on the task; however, similar results were obtained when we restricted the analysis to later blocks of the task (i.e., after participants have had more time to learn the environment).

### 2.2 Results

#### 2.2.1 Repetition Enhanced Pointing Performance and Increased Memory Strength

A linear regression model revealed an effect of block on performance on the JRD task, as assessed using median absolute angular error (henceforth, median error; see left panel of Figure 2). Specifically, a model that included block as a regressor fit the data better than a null model that only included a random effect of participant (i.e., random intercepts and slopes; block model AIC = 801; null model AIC = 822; likelihood ratio test: *X*^2^(1, *N* = 16) = 22.80, *p* < 0.0001). The *β* estimate for block was −7.04 (*t* = −6.86), suggesting that performance improves approximately 7 degrees per block. This finding replicates previous findings, showing that repetition, in the form of navigation, improves performance on the JRD task (Zhang et al., 2014). Similarly, a linear regression model revealed an effect of block on confidence ratings (*β* = 0.29, *t* = 8.28; block model AIC = 152; null model AIC = 177; likelihood ratio test: *X*^2^(1,*N* = 16) = 26.63, *p* < 0.0001; see middle panel of Figure 2), thus suggesting that confidence increases with repeated exposure to the environment.

**Figure 2:**
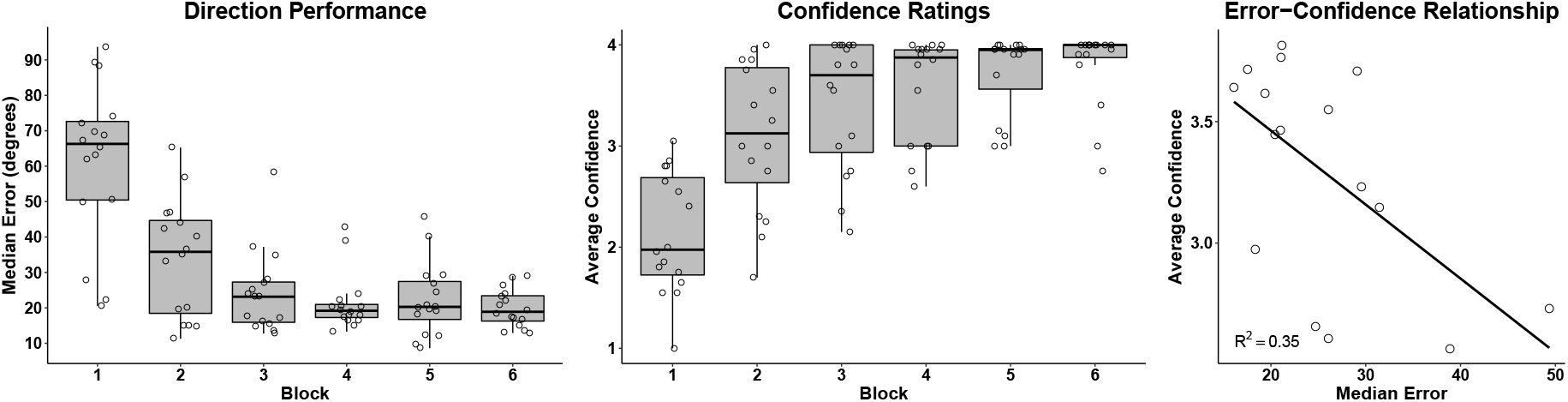
Performance on the JRD Task in Experiment 1. Repetition enhanced pointing accuracy (see boxplot in Left Panel) and subjective confidence (see boxplot in Middle Panel). There was a significant correlation between pointing error and confidence on the JRD task (see scatterplot in Right Panel).

We next investigated the relationship between confidence ratings and the JRD task and we observed a significant correlation. We found that higher confidence ratings correlated with lower pointing error (Pearson’s *r* = −0.59, *R*^2^ = 0.35, *t*(15) = −2.74, *p* = 0.016; see right panel of Figure 2). We also used an ordinal logistic regression model (using the clmm function within R package *ordinal*, version 2018.4-19) to investigate whether there was a relationship on a more within-trial basis—i.e., between confidence and pointing error across trials. Briefly, the full model used the angular error from all trials of the task across all participants to fit the confidence ratings data, and this model included a random intercept term across subjects (i.e., the trialwise data was fit across all subjects simultaneously). The full model fit the confidence ratings data better than a null model that only included random intercepts across participants (*β* = −0.021, full model AIC = 3506.0; null model AIC = 3789.9; likelihood ratio test: *X*^2^(1, *N* = 16) = 285.84, *p* < 0.0001). Together, these findings support the hypothesis that participants are aware, to some extent, of the accuracy of their spatial knowledge when responding in the JRD task.

### 2.3 Discussion

The results of Experiment 1 provide evidence that pointing accuracy improves over blocks of the task, which could be driven either by repeated learning (i.e., navigation) or repeated testing (i.e., blocks of the JRD task; for evidence of repeated testing effects on spatial memory see: Carpenter & Pashler, 2007; Carpenter & Kelly, 2012). These results converge with a wealth of evidence suggesting that repetition improves memory performance more generally (Ebbinghaus, 1913; Hintzman, 1976). Regarding the effect of repetition on spatial memory, the results have been equivocal, with some evidence of improvement in pointing accuracy with repeated exposure (Schinazi et al., 2013; Zhang et al., 2014; Starrett, Stokes, Huffman, Ferrer, & Ekstrom, 2018), while other studies have found little to no change with repeated exposure (Ishikawa & Montello, 2006). In Experiment 2 we aim to replicate this finding, which would provide stronger support for the hypothesis that repetition enhances the precision of spatial memory, as assayed by the JRD task.

Our results also suggest that subjective confidence increases with repeated navigation and testing. Moreover, we observed a relationship between overall pointing accuracy and subjective confidence. Interestingly, few previous studies have investigated the relationship between accuracy and confidence on the JRD task. Notably, Stevens and Carlson (2016) also observed a relationship between pointing accuracy and confidence on the JRD task. Taken together, these results extend the finding that performance on pointing tasks tends to correlate with subjective measures of navigational ability (Kozlowski & Bryant, 1977; Bryant, 1982; Ishikawa & Montello, 2006) to the realm of individual differences in subjective and objective measures of spatial navigation ability within a single environment.

## 3 Experiment 2

In Experiment 2, we aimed to replicate the findings of Experiment 1 and to test two additional hypotheses, as mentioned in the Introduction. First, we investigated the test-retest reliability of participants’ responses on the JRD task. Second, we tested the hypothesis that the JRD task and map drawing recruit similar cognitive representations. To this end, we implemented both a group-level analysis and a within-subject analysis in which we investigated whether the pattern of errors were correlated between the JRD task and the map drawing task. If these two tasks recruit similar cognitive representations, then we expect to find both a significant group-level relationship (i.e., evidence of individual differences in spatial cognition performance) and a significant within-subject relationship of the pattern of errors exhibited across the two tasks. The latter finding would strengthen the notion that participants recruited a similarly distorted representation to solve both tasks.

### 3.1 Method

#### 3.1.1 Participants

Thirty-one participants were recruited from the UC Davis community. Participants consented to the procedures in accordance with the Institutional Review Board of UC Davis and received course credit for their participation. Participants were between 18 and 43 years old (mean = 21.6; 17 females, 14 males). One participant was excluded because they left the session prior to completing the map drawing task. Four participants were excluded because they failed to perform significantly better than chance (one-tailed *p* > 0.05, as assessed by a permutation test; see 2.1.6 Data Analysis); therefore, 26 participants were included in the main analyses.

#### 3.1.2 Procedure

The virtual environment and the procedure for Experiment 2 were the same as Experiment 1 with two differences. First, the final block of the JRD task consisted of 60 questions (3 repetitions of 20 questions), which allowed us to investigate the test-retest reliability of the JRD task. Second, after completing all 6 rounds of tasks, participants performed a map drawing task.

#### 3.1.3 Map Drawing Task

In Experiment 2, participants performed a map drawing task after completing all 6 rounds of navigation and JRD tasks. The task consisted of an overhead view of the environment, including the wall boundaries of the environment (Figure 1; Stokes, Kyle, & Ekstrom, 2015). At the onset of the task, all 10 stores were placed on the left-hand side of the environment, outside of the wall perimeter. Participants used the mouse to complete the task. Whenever the cursor hovered over a store, its name was shown in the upper left-hand corner of the screen. Participants clicked on the stores and dragged them into position. Participants were allowed to move the stores in any order, and they could adjust stores after moving them within the boundaries of the environment. Prior to beginning the task, participants were instructed to double check their completed maps before submitting their answer. When participants were confident in their map, they pressed “Return” to submit their answer. The task was programmed to only accept the answer after participants had moved all 10 stores within the boundaries of the environment.

#### 3.1.4 Data Analysis

We used the same analytical approach as in Experiment 1, with the addition of an analysis of test-retest reliability and of map drawing performance, as detailed below.

*Test-Retest Reliability of Responses in the JRD Task*. In Experiment 2, we assessed the test-retest reliability of participants’ responses. During the final block of the JRD task, each participant answered the same twenty questions three times. We calculated the circular correlation coefficient between responses (using the package *circular* version 0.4-93 in R). Note, the circular correlation coefficient uses the same formula as Pearson’s correlation coefficient but it involves a transformation using the sine function, which normalizes the data into a circular space (e.g., to normalize for the fact that responses of −179 degrees and 179 degrees are quite similar in circular space but very different in linear space). For each participant we calculated the circular correlation coefficient between responses for each of the three possible combinations of comparisons. Prior to group-level analysis, correlation coefficients were Fisher’s r-to-z transformed (z[r]) using the following equation:

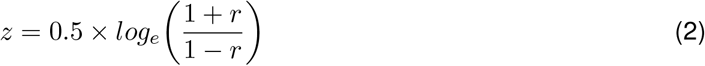

We then averaged the three z[r] circular correlation coefficients within each participant to obtain a mean coefficient for each participant. We assessed whether the relationship between responses was significantly better than zero using a one sample t-test. We also calculated the mean correlation coefficient for comparison to standard values of test-retest reliability. Specifically, mean correlation coefficients were calculated by applying the following equation to the mean z[r] correlation coefficients (Silver & Dunlap, 1987):

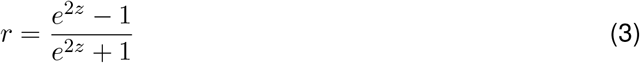

Silver and Dunlap (Silver & Dunlap, 1987) advocated for using backtransformed mean z[r] correlation coefficients (i.e., using Equations 2 and 3) rather than mean correlation coefficients (i.e., the mean of the raw correlation coefficients) because they are less biased estimates of the mean (as assessed by simulations). Thus, we adopted that approach here.

*Map Drawing Task*. We applied a metric transformation to each participant’s map using the package *nudged* (version 0.3.1; https://pypi.python.org/pypi/nudged) within Python (version 2.7.10). The metric transformation included translation, rotation, and uniform scaling. Uniform scaling preserves the angles between stores, which is motivated here by the fact that we used a square environment; furthermore, this method requires fewer assumptions than affine transformations that include non-uniform scaling, reflection, or shearing. To assess map drawing performance, we calculated the bidimensional regression coefficient between each participant’s metric-transformed map and the actual map coordinates using the R package *BiDimRegression* (version 1.0.6; Carbon, 2013; also see: Tobler, 1994; Friedman & Bernd, 2003). To assess whether there was a relationship between performance on the map drawing task and the JRD task, we calculated the correlation coefficient between performance on the map drawing task (as assessed by Fisher’s r-to-z transformed bidimensional regression coefficients) and the JRD task (as assessed by median error on the final block of the task).

To investigate whether the JRD task and the map drawing task recruit similar cognitive representations, we performed a trialwise analysis to investigate the consistency between errors on the JRD task and the map drawing task. Specifically, the signed error was calculated on the final 20 trials of the JRD task (i.e., the trials that took place immediately before the map drawing task) and the corresponding angles from the map drawing task. The signed error for the map drawing task was calculated in the same manner as for the JRD task—i.e., by comparing the actual angle to the angle drawn in the participant’s map for each of these JRD questions (see Left Panel of Figure 6). We then calculated the circular correlation coefficient (using the package *circular* in R) between the errors on the two tasks for each participant. As noted above, the circular correlation coefficient uses the same formula as Pearson’s correlation coefficient but it involves a transformation using the sine function. We then Fisher’s r-to-z transformed each participant’s circular correlation coefficient using Equation 2 and tested whether the group-level z[r] coefficients were significantly greater than 0; i.e., the null hypothesis was that there is no relationship between the errors on the JRD task and the map drawing task, which would be consistent with the hypothesis that these tasks recruit distinct cognitive representations.

### 3.2 Results

#### 3.2.1 Replication of Findings Related to Block Experiment 1

The results of Experiment 1 were replicated in Experiment 2. First, linear regression models revealed an effect of block on pointing performance (as assessed using median error; see left panel of Figure 3; *β* = −6.52, *t* = −8.67; block model AIC = 1339; null model AIC = 1372; likelihood ratio test: *X*^2^(1, *N* = 26) = 35.30, *p* < 0.0001) and on confidence ratings (see middle panel of Figure 3; *β* = 0.27, *t* = 7.50; block model AIC = 284; null model AIC = 312; likelihood ratio test: *X*^2^(1, *N* = 26) = 29.96, *p* < 0.0001) on the JRD task. Second, we observed a correlation between mean confidence and median error on the JRD task (Pearson’s *r* = −0.60, *R*^2^ = 0.35, *t*(24) = −3.63, *p* = 0.0013; see right panel of Figure 3). Furthermore, an ordinal logistic regression model that included angular error as a regressor fit the confidence ratings data better than a null model that only included random intercepts across participants (*β* = −0.017, full model AIC = 8421.3; null model AIC = 8858.9; likelihood ratio test: *X*^2^(1, *N* = 26) = 439.59, *p* < 0.0001).

**Figure 3:**
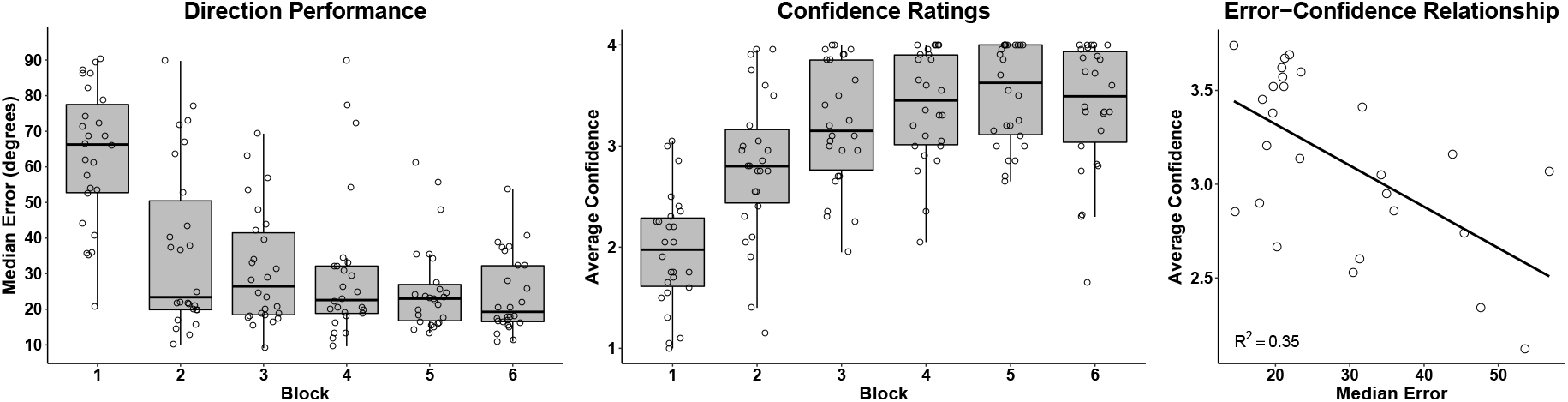
Performance on the JRD Task in Experiment 2. Repetition enhanced pointing accuracy (see boxplot in Left Panel) and subjective confidence (see boxplot in Middle Panel). There was a significant correlation between pointing error and confidence on the JRD task (see scatterplot in Right Panel).

#### 3.2.2 The JRD Task Exhibited a High Degree of Test-Retest Reliability

In Experiment 2, we assessed the test-retest reliability of the JRD task (see 3.1.4). The mean of the group z[r] circular correlation coefficients—i.e., our measure of test-retest reliability—was significantly greater than 0 (mean z[r] circular correlation coefficient =1.19, *t*(25) = 8.50, *p* < 0.0001; see left panel of Figure 4). Moreover, the magnitude of the effect suggests that there was a high degree of test-retest reliability; specifically, the mean circular correlation coefficient was 0.83 (calculated using Equation 3) and the median circular correlation coefficient was 0.85 (i.e., these values suggest that the relationship between responses across repetitions accounts for a large percentage of the variance in responses). We acknowledge that the magnitude of these relationships could be inflated due to the nature of our test (i.e., these questions occurred in the same block); however, designs that compare responses across different blocks would be subject to different assumptions that might not be met (e.g., responses might change as a result of learning or interference during delayed testing). Thus, our results provide support for the test-retest reliability of the JRD task.

**Figure 4:**
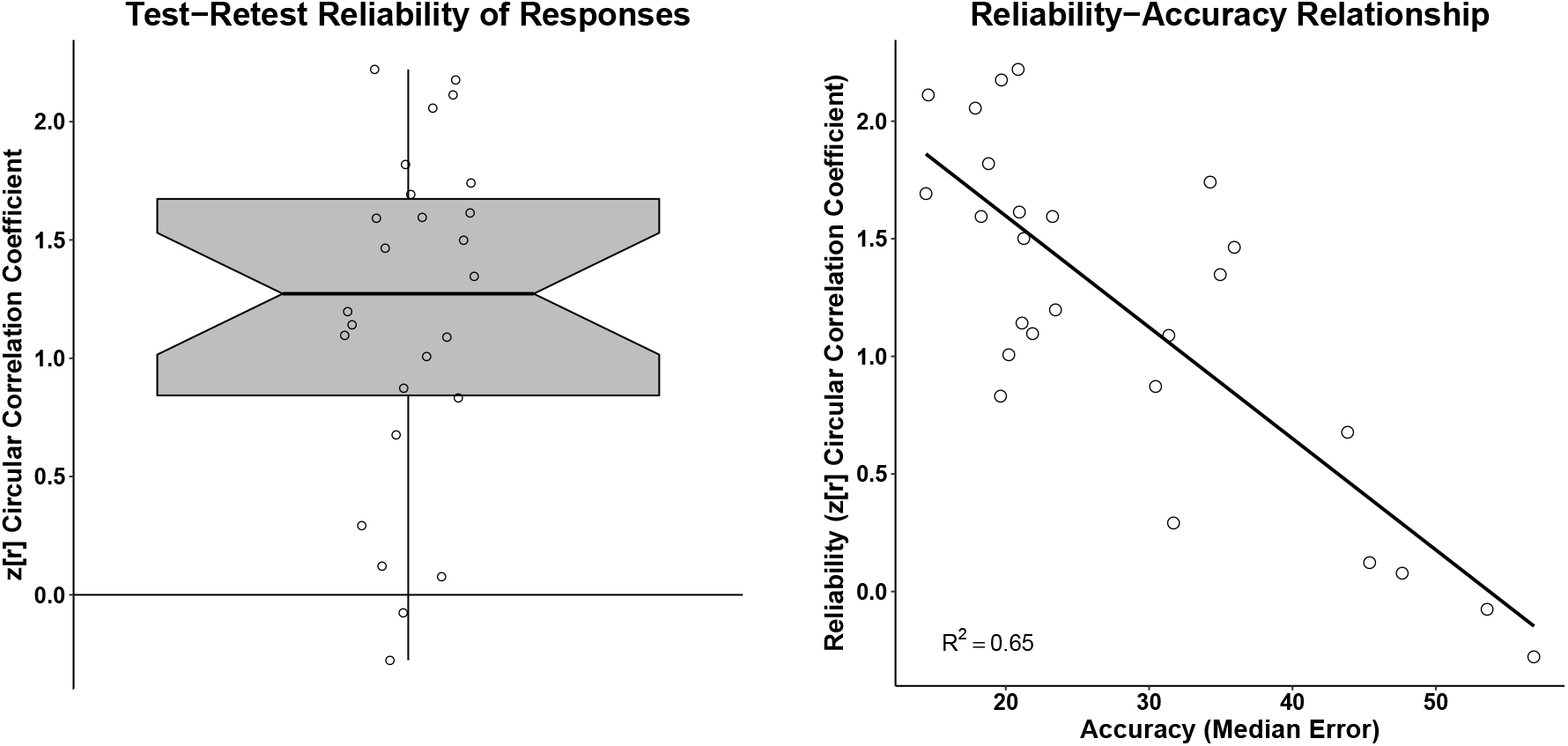
The JRD Task exhibits a high degree of test-retest reliability. The mean of the group z[r] circular correlation coefficients was significantly greater than 0 (mean z[r] circular correlation coefficient = 1.19, mean circular correlation coefficient = 0.83, median circular correlation coefficient = 0.85; see the notched boxplot in Left Panel). There was a significant correlation between the reliability of responses (mean z[r] circular correlation coefficient) and accuracy (median error across all blocks of the task; see scatter plot in Right Panel), suggesting that participants that are more consistent with their responses tend to also perform better on the task.

While the group exhibited a high degree of test-retest reliability overall, there was also a relatively broad range of test-retest reliability scores and we were interested in testing whether this variability was related to performance on the task (i.e., angular error). We observed a relationship between the reliability of responses (mean z[r] circular correlation coefficient) and accuracy (median error across all blocks of the task; Pearson’s *r* = −0.81, *R*^2^ = 0.65, *t*(24) = −6.71, *p* < 0.0001; see right panel of Figure 4). Altogether, these results suggest that the JRD task exhibits good test-retest reliability and participants that are more consistent with their responses—i.e., across the three repetitions of the same questions—tend to also perform better on the task.

#### 3.2.3 Correlation between JRD Task Performance and Map Drawing Accuracy

We next investigated whether there was a relationship between JRD task performance and map drawing, which is often considered a relatively pure measure of an allocentric representation. If the two tasks rely on similar cognitive processes, then we expect to observe a correlation between performance on the two tasks. If, however, these tasks recruit distinct cognitive processes, then we should not observe a significant relationship between performance on the two tasks. We first aligned each participants map data using a metric transformation to deal with the issue that different participants may have different preferred orientations at which they draw their map (see 3.1.4; all participant’s metric-transformed data can be seen in Figure 5). Next, we calculated the bidimensional regression coefficient of these metric-transformed maps, which revealed that participants performed quite well on the map drawing task overall (mean *r* = 0.96, mean *R*^2^ = 0.93; note, at the individual subject level, *p* < 0.0001 for 24 of 26 participants, *p* = 0.013 for one participant, and *p* = 0.12 for one participant). We observed a relationship between performance on the map drawing task (assessed via Fisher’s r-to-z transformed bidimensional regression coefficients) and performance on the JRD task (assessed via median error on the final block; Pearson’s *r* = −0.68, *t*(24) = −4.58, *p* = 0.00012; see Figure 5). Note, the between-task relationship maintained when two potential outliers (i.e., those with map performance scores < 1) were removed from the analysis (Pearson’s *r* = −0.43, *t*(22) = −2.26, *p* = 0.034). Additionally, similar results were obtained when using nonparametric rank correlation techniques (Spearman’s *ρ* = −0.65, *p* = 0.00048; Kendall’s *τ* = −0.50, *p* = 0.00025), which are less sensitive to potential outliers than Pearson’s correlation, and these results also maintained when removing the two potential outliers (Spear-man’s *ρ* = −0.55, *p* = 0.0059; Kendall’s *τ* = −0.41, *p* = 0.0042). Taken together, these results suggest that the group-level relationship between performance on the JRD task and the map drawing task is stable and is driven neither by the presence of outliers nor by the assumption of a linear relationship between performance on the two tasks (i.e., the nonparametric correlation approaches assume a monotonic but not necessarily linear relationship). Given that we were able to explain approximately 47% of the variance in JRD performance with map drawing, our findings suggest that these tasks tap into similar cognitive processes.

**Figure 5:**
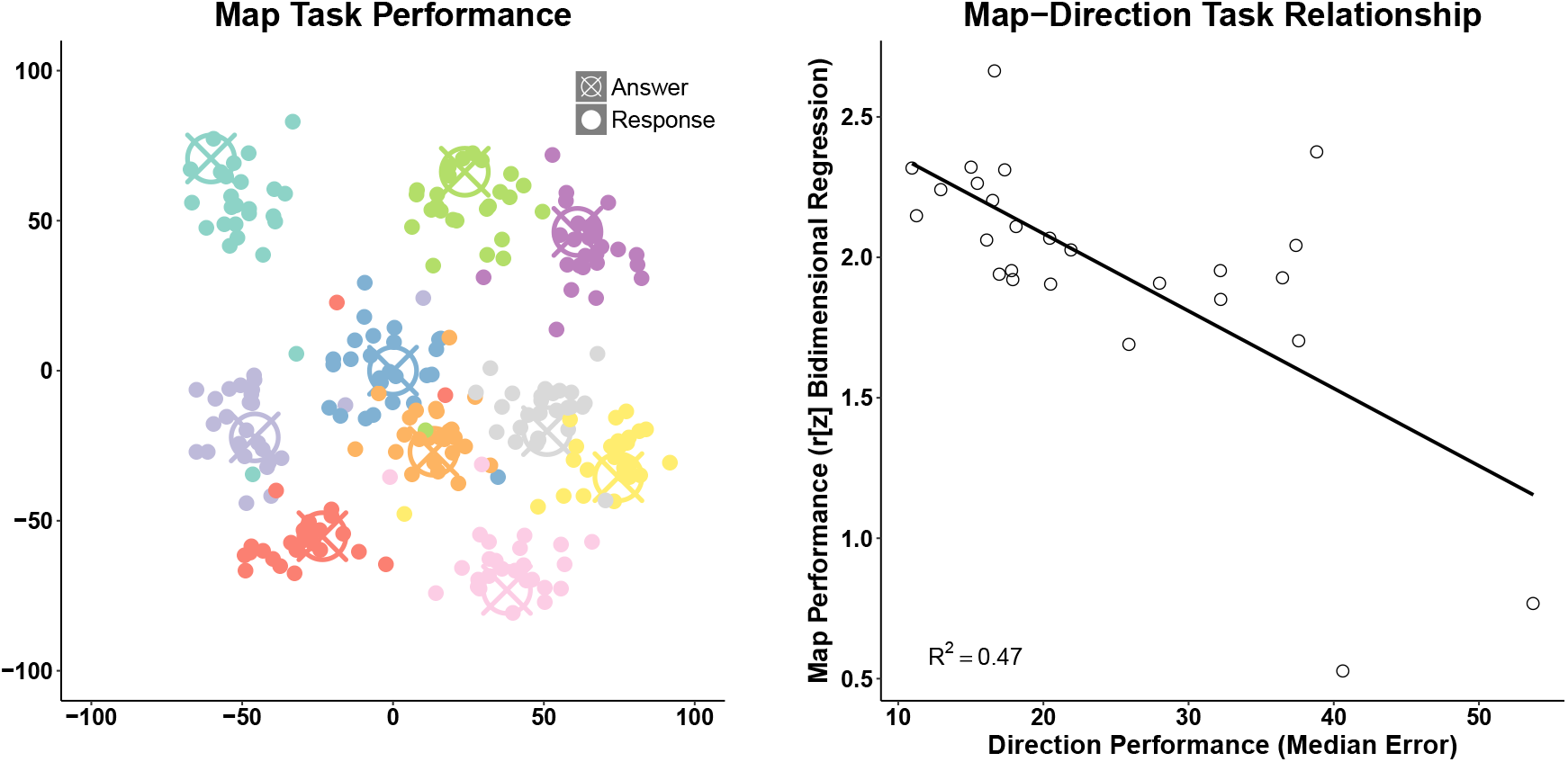
Performance on the map drawing task in Experiment 2. Participants performed well on the map drawing task overall (see the metric-transformed map data from all participants in Left Panel). There was a significant correlation between performance on the map drawing task (assessed via z[r] bidimensional regression coefficients) and performance on the JRD task (assessed via median error on the final block; see scatter plot in Right Panel).

#### 3.2.4 Similar Pattern of Angular Errors on the JRD Task and the Map Drawing Task

The results from the previous section suggest that there is a relationship between how well participants perform on the JRD and map drawing tasks. While interesting in their own right (e.g., from a psychometrics standpoint), the results of a group-level analysis can only provide indirect evidence that participants recruited similar cognitive representations to perform the two tasks. For example, it could be that there is a “global” spatial ability, and those that have this simply perform better between any spatial task, including the JRD and map drawing tasks (Hegarty et al., 2006). Thus, we more directly addressed this question by testing whether performance on the JRD and map drawing tasks were subject to similar distortions using a within-subjects trialwise analysis (see 3.1.4 and left panel of Figure 6), which would in turn provide more direct evidence that the two tasks tap into overlapping forms of representations.

**Figure 6:**
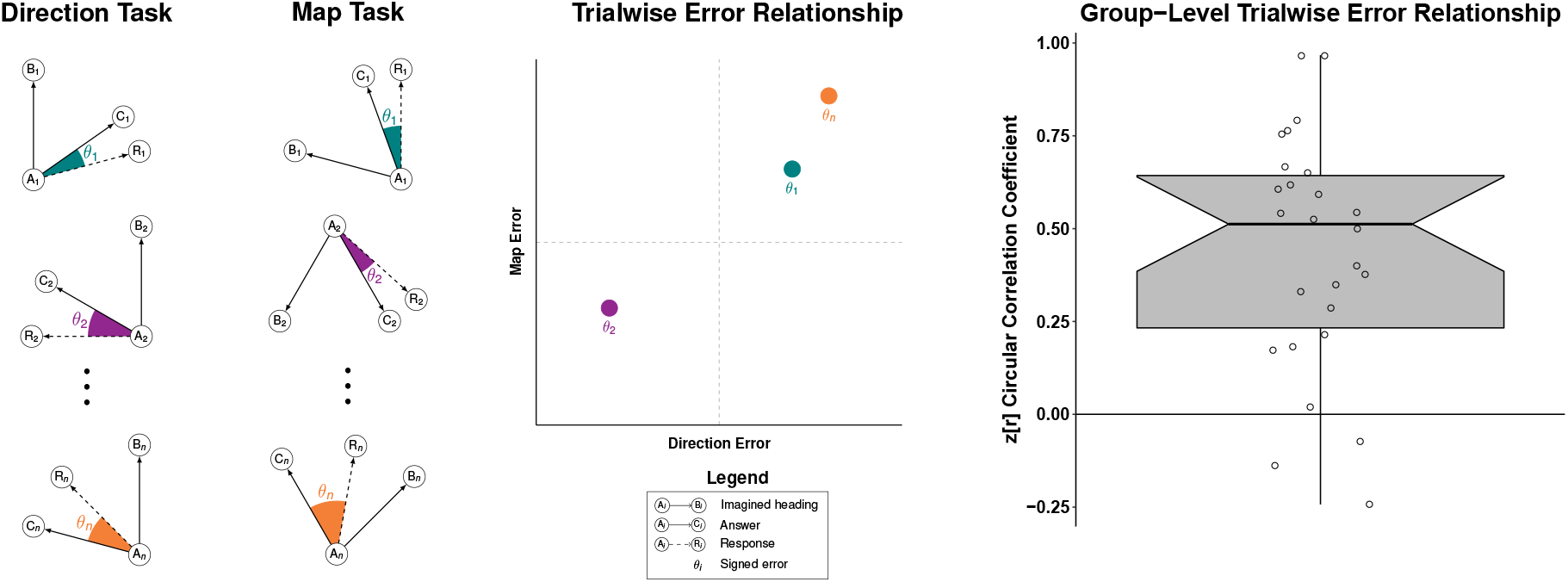
Trialwise analysis of the pattern of errors on the JRD task and the map drawing task. The signed error was calculated on each trial of the JRD task and the map drawing task (see Left Panels). We then calculated the circular correlation coefficient between the errors on the two tasks for each participant (see Middle Panel). The trialwise error relationship was significantly greater than zero (see notched box plot in Right Panel; each dot represents an individual participant; note, notched box plots give an estimate of the 95% confidence interval of the median, and the lower notch in this boxplot does not include 0, thus providing converging evidence that there is a significant relationship between the trialwise pattern of errors on the two tasks).

We hypothesized that if performance on the two tasks rely on similar cognitive representations, then errors in a participant’s judgment of the angles between stores should be correlated between the two tasks. Conversely, if the JRD and map drawing tasks rely on distinct cognitive representations, then we should fail to observe a relationship between the trialwise error on the two tasks. Consistent with our hypothesis, we observed a significant relationship between the trialwise angular error on the JRD and map drawing tasks (mean z[r] circular correlation coefficient = 0.44, *t*(25) = 6.95, *p* < 0.0001; Figure 6). We would like to emphasize that the trialwise analysis assessed the relationship between the errors on the two tasks, as opposed to the overall relationship between the angles across the two tasks. The latter approach would be correlated with the true angles between the stores; in fact, the correlation was higher for the overall relationship (median circular correlation coefficient = 0.84) than the trialwise error relationship (median circular correlation coefficient = 0.47). Therefore, our analysis of trialwise errors provides a cleaner measure of the similarity of the distortions in participants’ cognitive representations.

### 3.3 Discussion

Our first relevant finding here is that the JRD task shows good test-retest reliability. Interestingly, we also observed a strong relationship (*R*^2^ = 0.65) between participants’ performance on the task and their test-retest reliability. These results suggest that participants that tend to perform better on the task tend to also exhibit more reliable responses to the same questions across repeated testing. These results are important because they support the construct validity of the JRD task. Moreover, these results suggest that better performing participants are both more accurate and less variable in their responses. Thus, poorer performing participants might employ more variable cognitive representations relative to better performing participants. This hypothesis could be further tested by investigating the relationship between task performance and variability of neural representations of spatial information, for example, by applying multivariate pattern analysis to fMRI or EEG data.

We presented two pieces of evidence that the JRD task and the map drawing task recruit similar cognitive processes. First, we found a significant correlation between performance on the two tasks, supporting previous studies that have found a correlation between performance on the map drawing task and both the SOP task (Hegarty et al., 2006; Labate et al., 2014; Weisberg et al., 2014; but see: Schinazi et al., 2013) and the JRD task (Bryant, 1984; but see: Schinazi et al., 2013). Thus, our results add to a growing body of evidence to suggest that there is a relationship between performance on pointing tasks, such as the JRD task, and map drawing tasks. As mentioned in the Introduction, however, a more direct analysis is necessary to conclude that participants recruit similar cognitive representations to solve the JRD and map drawing tasks (cf. Bryant, 1984), which we addressed in our second analysis.

We found a significant within-subject relationship between the trialwise angular errors on the JRD task and the map drawing task. Because this analysis assessed the between-task correspondence of errors within individual participants on a trial-by-trial basis, these results provide more direct support for the notion that participants recruit similarly distorted cognitive representations to solve the two tasks. Moreover, these results suggest that the JRD task provides a useful measure of holistic representations of the environment. A previous study reported a more modest relationship between the pattern of errors on the two tasks (Bryant, 1984). However, there are important differences between the analyses used in these two studies. For example, the previous study investigated the similarity of error patterns using a group-level analysis. In contrast, our analysis was conducted within subject, thus allowing a more direct estimate of systematic distortions of cognitive representations within individual participants (i.e., something akin to a participant-specific “cognitive map”). Additionally, the map drawing task in Bryant (1984) provided participants with a detailed map of the environment overlaid on a grid of squares, and participants were instructed to write the number of the corresponding grid square next to the name of each landmark, which was written outside of the map. Together, these differences may have deemphasized participants’ angle estimates between landmarks. In contrast, our map drawing task encouraged participants to consider the relative locations of all stores simultaneously—i.e., it is likely that participants in our task paid greater attention to the relative angles and distances between stores, as opposed to focusing on each store in relative isolation. Finally, Bryant (1984) used a real-world environment rather than a virtual environment. Taken together, the results of our group-level and within-subjects analyses suggest that the JRD task and the map drawing tasks recruit similar cognitive representations, and future studies could investigate whether there are conditions in which this relationship is enhanced or diminished.

## 4 Simulation of Chance Performance on the JRD Task

As mentioned in the Introduction, one question regarding the JRD task is whether or not chance performance depends on the distribution of participants’ responses. To address this question, we simulated performance on the JRD task to investigate the effects of various parameters, including the distribution of the angles of responses, on performance. If chance performance is stable and 90 degrees, as is commonly assumed (Sholl, 1987; Werner & Schmidt, 1999; McNamara, Rump, & Werner, 2003; Waller & Hodgson, 2006; Tlauka, Carter, Mahlberg, & Wilson, 2011; Zhang et al., 2014; Street & Wang, 2016), then our simulations should reveal that chance performance is 90 degrees regardless of the distribution of responses. We note that the answer to this question depends somewhat on the experimental design. We return to this issue in the General Discussion.

### 4.1 Method

We used the store locations from our virtual environment in the simulations—i.e., the answers to each question were based on the locations in the virtual environment. Each simulated participant was asked every unique question across the 10 stores (720 questions; *n* × (*n* − 1) × (*n* − 2), where n is the number of target stores). We simulated responses by drawing from uniform distributions with constraints on the lower and upper bounds of the distribution. Specifically, for each set of parameters, we simulated 720 responses for each simulated participant and calculated the absolute angular error between these simulated responses and the actual angles. To estimate the mean angular error^3^, we simulated 10,000 participants for each set of parameters and we calculated the mean angular error for each participant.

### 4.2 Results

When simulated responses were drawn from a uniform distribution between 0 and 360 degrees (minimum angle, maximum angle, respectively), the resulting mean angular error (average of the 10,000 participant-specific averages) was approximately 90 degrees (see orange distribution in Figure 7), as is commonly assumed. In contrast, the mean angular error was less than 90 degrees whenever the distribution of responses was constrained toward responses in front of the simulated participant. For example, when responses were constrained to the forward-facing 180 degrees (i.e., between −90 and 90 degrees), the mean error was substantially less than 90 degrees (see purple distribution), and constraining the responses between −45 and 45 degrees further shifted the null distribution away from 90 degrees and closer to 60 degrees (see green distribution). The reason that mean angular error decreases when responses are constrained toward forward-facing responses is due to an effect that we refer to as “the triangle rule.” Specifically, because each triad of stores forms a triangle, the mean absolute angle across all such triangles in the city is exactly 60 degrees (i.e., 180/3; see left panel of Figure 7). Thus, when every response (across all possible questions) is made with the pointer facing directly forward (i.e., response of 0 degrees), the mean absolute angular error is always exactly 60 degrees (see dashed line). These results demonstrate the mean angular error depends on the distribution of the angles of responses.

**Figure 7:**
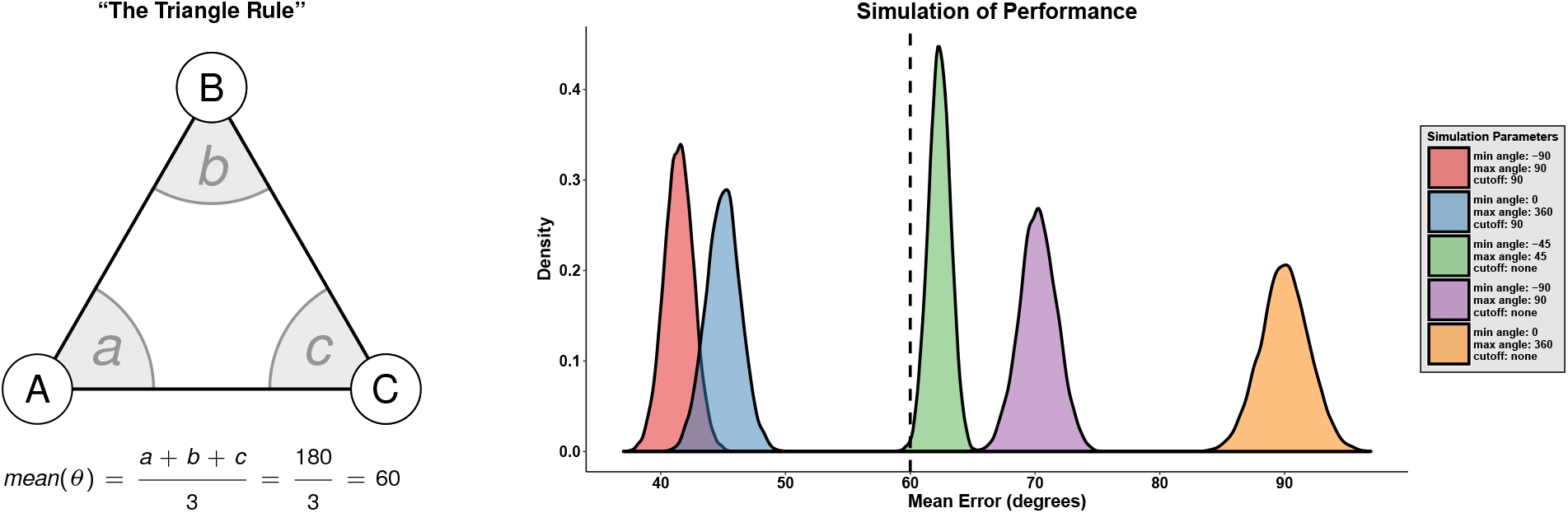
Simulation of chance performance on the judgment of relative directions task. The average absolute angle for each triad of questions (i.e., all permutations of the following question, “Imagine you’re standing at A, facing B. Please point to C.”) is 60 degrees due to “the triangle rule” (Left Panel). The triangle rule causes the chance angular error, rather than being stable and 90 degrees, to differ based on the distribution of the directions of simulated responses (see the orange, purple, and green distributions in Right Panel). Other analytical decisions, such as removing trials that exceed a maximum angular error cutoff, also shift chance errors to be less than 90 degrees (see the blue distribution) and the distribution shifts further to left when the distribution of responses is also constrained in the forward-facing direction (see the red distribution). The dashed line represents the condition in which all responses are pointed directly in front of the simulated participant (the mean error for all of the simulated participants in this case was exactly 60 degrees due to the triangle rule).

Previous studies have implemented a maximum angular error cutoff to minimize the influence of outliers from comparisons of pointing accuracy (e.g., 90 degrees in Sholl, 1987; Werner & Schmidt, 1999; McNamara et al., 2003; Street & Wang, 2016). We were interested in studying the effect of this procedure, so we simulated the effect of removing errors that exceeded a given cutoff. Employing a cutoff of 90 degrees, at least based on our data derived from our experimental design, had an impact on the distribution of errors; specifically, the resultant mean error was half of the given cutoff when responses were simulated from a uniform distribution between 0 and 360 degrees. For example, with a cutoff of 90 degrees, the mean error was approximately 45 degrees (see blue distribution). Additionally, removing responses that exceeded a variability criterion (e.g., 2 standard deviations from the mean; Zhang et al., 2014) also reduced the mean angular error (not shown in figure), which was likely driven by the fact that the underlying error distributions are skewed, causing only large errors to be excluded by this procedure (i.e., small errors were never 2 standard deviations from the mean). These results indicate that removing trials that exceed a threshold, either via maximum error or standard deviations from the mean, can significantly influence chance performance (i.e., shift the null distribution to be less than 90 degrees).

We next investigated the effect of constraining simulated responses (e.g., between −90 and 90 degrees) with a maximum angular error cutoff (e.g., 90 degrees) and we observed a combined effect of these two parameters (see red distribution). This finding is particularly important because the actual distribution of the answers is not uniform around the circle (i.e., uniform from 0 to 360 degrees). Instead, the distribution of answers is biased toward forward-facing angles (i.e., due to the triangle rule). Thus, if the distribution of the responses is constrained to reflect the distribution of the answers, then the distributions will be shifted to be less than 90 degrees. Moreover, applying a maximum angular error cutoff additionally shifts the distributions further from 90 degrees.

### 4.3 Discussion

The results of our simulations demonstrate that chance performance differs based on the distribution of responses, which in turn can differ based on how a participant distributes their guesses. Thus, our simulations suggest that chance performance on the JRD task is not consistently 90 degrees and instead depends on a bias toward a forward-facing angle due to the triangle rule (i.e., forward-facing responses result in an average pointing error of 60 degrees, the average angle of a triangle). Instead, when tests versus chance performance are conducted, we advocate for the use of methods that take the distribution of responses into account (e.g., permutation tests). In the next section, we will demonstrate an example of a group-level permutation approach for testing whether performance differs from chance at the group level (see Group-Level Permutation Test). For example, the group-level permutation test can be useful for determining whether a population performs better than chance on the JRD task (e.g., healthy older adults, patients with memory-related disorders such as mild cognitive impairment or Alzheimer’s disease).

In addition to the effect of the distribution of participants’ responses on chance performance, outlier removal can have a dramatic influence and this effect is further pronounced when the distribution of responses is biased toward forward-facing angles. One alternative approach for mitigating the influence of outliers in the data (within participants) is to use the median error because it is less influenced by the presence of outliers than the mean error—i.e., the within-subject angular error distributions are skewed and outliers add to this skewed positive tail, thus biasing within-subject means more than medians. In fact, we used the median error (within participants) for analyzing our empirical data. Additionally, for one-sample experiments, we advocate for participant exclusion based on whether or not they performed better than chance on the task overall (see 2.1.6 Data Analysis). For example, participant exclusion is particularly helpful for testing for within-subjects differences in performance due to differences in an experimentally defined independent variable because participants that perform at a chance level overall will only contribute noise to such statistical effects (e.g., due to floor performance across conditions). Permutation tests will also potentially exclude participants that appear to perform better than chance as assessed by measures of angular error alone. For example, participants that always point directly in front of themselves will have a mean angular error of 60 degrees but will perform at chance level (e.g., p = 1). In contrast, it is possible that participants will perform with a mean angular error that appears to be “worse” (e.g., 70 degrees), but due to the distribution of their responses will perform better than chance. Thus, permutation tests provide a straightforward assessment of whether a participant performs better than chance and the use of such tests can remove noisy participants from a within-subjects analysis. Note, however, that participant exclusion should not be conducted in cases in which overall performance is compared to chance at the group level (e.g., as mentioned above in patients with memory-related disorders) because it will be guaranteed to be better than chance (i.e., because the participants performing at chance will have already been excluded). Finally, these results provide an indirect example of the importance of matching the distribution of the angles of answers when comparing performance between different conditions (cf. Montello, Richardson, Hegarty, & Provenza, 1999), as has been done in previous studies (e.g., Mou, McNamara, Valiquette, & Rump, 2004; Mou, Zhao, & McNamara, 2007; Shelton & Marchette, 2010).

In summary, we advocate for the use of median error (within subject) as opposed to removing “outlier” trials. Additionally, participant exclusion can be helpful when conducting a within-subjects analysis of performance on the JRD task. In the next two sections, we provide empirical evidence that corroborates the findings of our simulations.

## 5 Empirical Investigation of Guessing Performance on the JRD Task

We next asked whether guessing performance also differed from 90 degrees in our empirical data using confidence ratings. Specifically, we calculated the mean angular error for “guessing” trials (i.e., trials with confidence ratings of 1=“very unsure”). Note, for comparison to other studies, we calculated the mean angular error in this analysis—as discussed in 4.3, for all other analyses, we calculated the median angular error as a measure of performance within each participant because it is less influenced by outliers than mean angular error. If chance performance on the JRD task is 90 degrees, then we would expect to observe a mean error for guessing trials that is not significantly different than 90 degrees. Conversely, if the mean error for guessing trials is significantly better than 90 degrees, as suggested by the results of our simulations, then it would suggest that chance performance is better than 90 degrees.

### 5.1 Method

#### 5.1.1 Participants

We included the data from all participants from Experiments 1 and 2.

#### 5.1.2 Group-Level Permutation Test

We used a group-level permutation test to investigate whether group-level performance differed from chance. For this procedure, we used two-tailed nonparametric p values, which were calculated using the following equation (Ernst, 2004; Huffman & Stark, 2014, 2017):

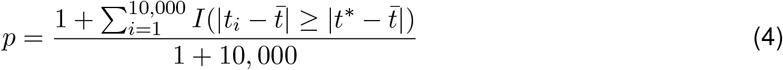

where 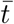 is the mean of the null distribution. We derived the group-level null distribution using a two step procedure. First, we ran 10,000 permutations within each participant. For each permutation, we randomly shuffled the participant’s responses and calculated the mean error with these arbitrarily associated correct angles (i.e., the result of this step was *N* vectors of 10,000 elements). Second, we constructed a group-level null distribution by averaging these vectors across all of our participants (i.e., resulting in a vector of 10,000 elements, each element of which is a randomized average performance across participants). This approach is analagous to permutation tests used in previous neuroimaging analyses (e.g., Chen et al., 2011; Liang, Mouraux, Hu, & Iannetti, 2013; Stelzer, Chen, & Turner, 2013; Etzel, 2015; Huffman & Stark, 2017).

We also ran another procedure in which we compared each participant’s empirical mean absolute angular error to the mean of their null distribution using a paired t-test approach^4^, which provided another technique for controlling for the differences in null pointing error across participants (e.g., due to putative differences in response biases). Additionally, to assess the evidence for or against the null hypothesis, we also implemented a Bayes factor version of this analysis using the *ttestBF* function from the package *BayesFactor* (0.9.12) in R (with the *rscale* parameter set to the default of 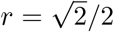; Rouder, Speckman, Sun, Morey, & Iverson, 2009; Morey & Rouder, 2011).

### 5.2 Results

To increase statistical power, we included all participants from both Experiment 1 and 2, whether or not they performed better than chance on the task, with the condition that they had at least 10 trials with confidence ratings of 1=“very unsure” (*N* = 25). We then calculated the mean error for each participant and tested whether the mean errors differed from 90 degrees using a one-sample t test. We found that the mean error on guessing trials was significantly better than 90 degrees (mean error = 81.8, *t*(24) = −2.76, *p* = 0.011) and a Bayes factor t-test revealed evidence in favor of the alternative hypothesis (*BF*_10_ = 4.4, i.e., the alternative hypothesis is approximately 4.4 times more likely than the null hypothesis; Figure 8). These results could either suggest that: 1) participants’ guesses contain above-chance information, or 2) the null distribution is shifted to be less than 90 degrees. We used two permutation tests to adjudicate between these two possibilities (see Group-Level Permutation Test), and we found that performance on guessing trials did not differ significantly from chance using our group-level permutation approach (*p* = 0.34) and our group-level paired t-test approach (mean difference = −2.1, *t*_24_ = −0.75, *p* = 0.46, Bayes factor in favor of the null [*BF*_01_] = 3.7; Figure 8). In contrast, performance was better than chance for all of the other confidence ratings under both methods (all *p* < 0.0001; all *BF*_10_ > 11,000).

**Figure 8:**
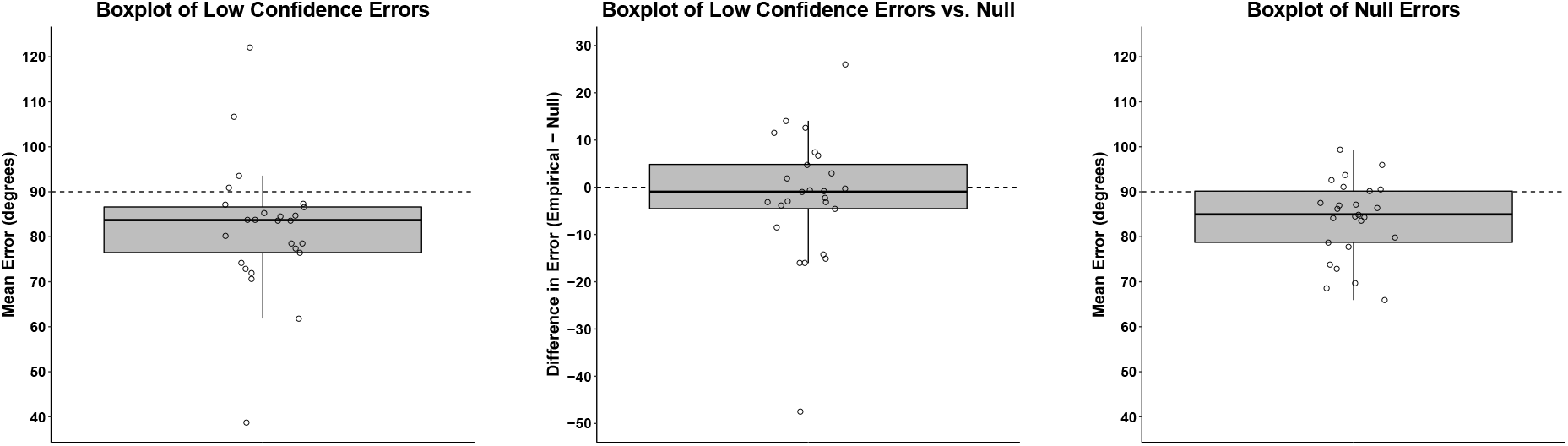
Performance on guessing trials revealed the importance of using permutation tests for assessing chance performance on the JRD task. A parametric analysis revealed that the mean error on guessing trials (i.e., trials in which confidence ratings were 1=“very unsure”) was significantly better than 90 degrees (see boxplot in Left Panel). In contrast, our permutation tests revealed that performance on guessing trials was not significantly different than the permutation-derived chance performance (see boxplot in Middle Panel). Moreover, 90 degrees was significantly better than the mean error across all participant’s null distributions (see boxplot in Right Panel; note: each dot represents a single subject’s mean of their null distribution).

Next, we assessed whether 90 degrees error was significantly worse than the permutation-derived null distributions from both our group-level permutation approach and our paired t-test approach. We found that 90 degrees was significantly worse than the mean of the group-level null distribution (*p* < 0.005), and this result was replicated by the group-level null distributions for confidence ratings 2=“somewhat unsure” (*N* = 40, *p* < 0.0001), 3=“somewhat sure” (*N* = 48, *p* < 0.0001), and 4=“very sure” (*N* = 40, *p* < 0.0001). Finally, these results were replicated by our paired t-test approach. Specifically, we conducted a Bayes factor t-test between the group of permutation-derived null angular errors (i.e., each value was the mean of a subject’s permutation-derived null distribution) and we found that these null values were associated with significantly less angular error than 90 degrees for confidence ratings 1=“very unsure” (mean = 83.9; *t*_24_ = −3.54, *p* = 0.0017, *BF*_10_ = 22.1; Figure 8), 2=“somewhat unsure” (mean = 82.0, *t*_39_ = −5.67, *p* < 0.0001, *BF*_10_ = 11,664), 3=“somewhat sure” (mean = 80.8, *t*_47_ = −8.90, *p* < 0.0001; *BF*_10_ = 769,441,659), and 4=“very sure” (mean = 77.2, *t*_39_ = −8.86, *p* < 0.0001, *BF*_10_ = 143,713,465). Altogether, our findings including confidence ratings again suggest that chance performance on the JRD task is not consistently 90 degrees.

### 5.3 Discussion

Consistent with our simulations, our empirical data revealed that guessing trials are associated with a mean angular error that is significantly less than 90 degrees, as assessed via parametric assumptions of chance performance of 90 degrees. However, our group-level nonparametric permutation method suggested that guessing performance does not differ significantly from chance. Additionally, we found that 90 degrees is, in fact, significantly worse than the group-level null distributions across all levels of confidence. Together, these empirical results challenge the assumption that chance performance is 90 degrees, and we suggest that nonparametric permutation tests should be used to derive null distributions without assuming chance performance of 90 degrees. Specifically, when testing whether a group of participants perform better than chance on the JRD task, we advocate for the use of a group-level permutation approach (see Group-Level Permutation Test) as opposed to making parametric assumptions with chance performance of 90 degrees.

## 6. Experiment 3

In Experiment 3, we aimed to extend the results from our guessing analysis (see Empirical Investigation of Guessing Performance on the JRD Task) to a state in which participants truly had no knowledge of the environment. Thus, we had participants perform the JRD task without exposing them to the environment. If performance under such conditions is still significantly better than 90 degrees, then it would further support the conclusion that chance performance can differ from 90 degrees.

### 6.1 Method

#### 6.1.1 Participants

Twenty-five participants—i.e., the number of participants that met criteria for inclusion in the low-confidence ratings data—were recruited from the UC Davis community. Participants consented to the procedures in accordance with the Institutional Review Board of UC Davis and received course credit for their participation. Participants were between 18 and 26 years old (mean = 20.0; 22 females, 3 males).

#### 6.1.2 Procedure

Participants performed 60 trials of the JRD task without being exposed to the virtual environment. Participants were given the following instructions: “Our lab studies navigation and spatial memory. Thus, we typically have participants navigate in virtual environments (i.e., environments that exist on the computer but not in the real world). We test participants’ memory for these environments using the task that you will be performing today, namely, a pointing task. You are in a control condition in which you will not actually navigate the city. Thus, your task today will be to *GUESS* where stores are located in the virtual environment.”

### 6.2 Results and Discussion

Mean absolute angular error was significantly better than 90 degrees in the no-navigation group (mean error = 83.8 degrees, *t*_24_ = −4.12, *p* = 0.00039, *BF*_10_ = 80.3; Figure 9), similar to the results of our low-confidence ratings data. In contrast, and as expected, both our group-level permutation method (*p* = 0.20) and our paired t-test approach (mean difference = −1.6, *t*_24_ = −1.33, *p* = 0.20, *BF*_01_ = 2.2) correctly failed to reject the null hypothesis of chance performance in the non-exposed group (Figure 9). Moreover, 90 degrees was again significantly worse than the mean error of the null distribution from our permutation analysis (*p* < 0.0001) and similar results were obtained when we assessed whether the mean of each participant’s null distribution was less than 90 degrees (mean = 85.4, *t*_24_ = −4.43, *p* = 0.00018, *BF*_10_ = 160.1; Figure 9).

**Figure 9:**
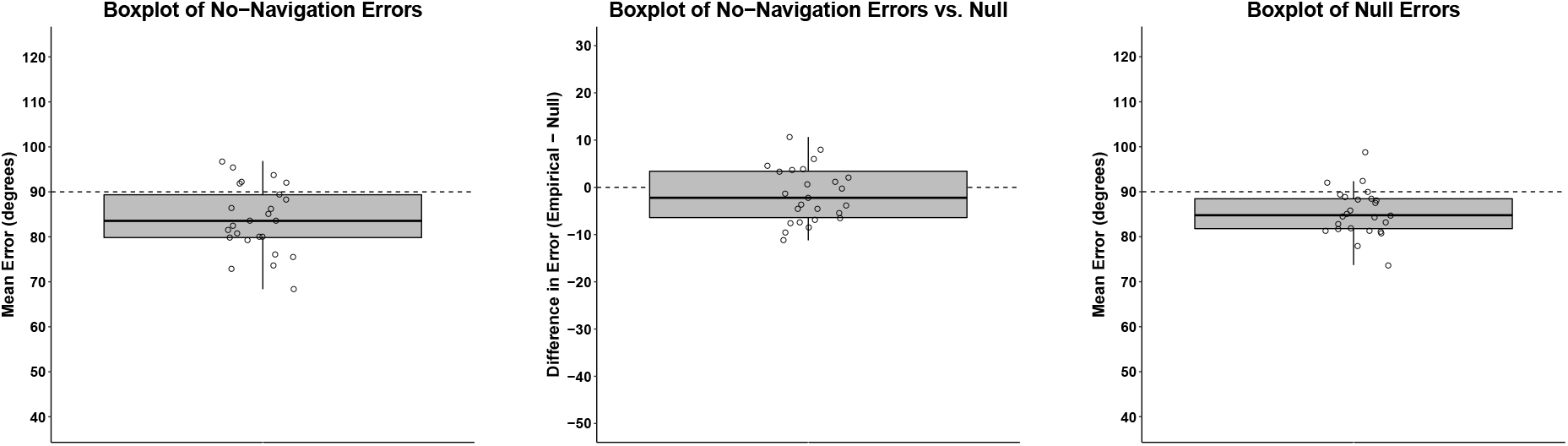
Performance in the no-navigation group revealed the importance of using permutation tests for assessing chance performance on the JRD task. A parametric analysis revealed that the mean error in the no-navigation group (i.e., participants that never navigated the virtual environment) was significantly better than 90 degrees (see boxplot in Left Panel). In contrast, our permutation tests revealed that performance in the no-navigation group was not significantly different than the permutation-derived chance performance (see boxplot in Middle Panel). Moreover, 90 degrees was significantly better than the mean error across all participant’s null distributions (see boxplot in Right Panel; note: each dot represents a single subject’s mean of their null distribution).

Next, we compared performance between the no-navigation group and the confidence ratings 1=“very unsure” group. A two-sample t-test revealed that performance in the low-confidence ratings group and the non-exposed group did not significantly differ (*t*_48_ = −0.60, *p* = 0.55, *BF*_01_ = 3.04). When we combined the data from both of these groups, we again found that pointing error was significantly better than 90 degrees (mean = 82.8, *t*_49_ = −4.35, *p* < 0.0001, *BF*_10_ = 321), and our paired t-test Bayes factor analysis revealed evidence in favor of the null hypothesis of no difference from chance (mean difference = −1.8, *t*_49_ = −1.22, *p* = 0.23, *BF*_01_ = 3.3). Altogether, the results from our simulations, the low-confidence ratings group, the no-navigation group suggest that chance pointing accuracy can deviate from 90 degrees (at least under the condition in which questions are sampled randomly from all possible questions).

## 7 General Discussion

### 7.1 Construct Validity of the JRD Task

Previous studies have investigated the construct validity of pointing tasks (Montello et al., 1999; Waller, Beall, & Loomis, 2004), but these tasks have not focused specifically on the JRD task. These studies also focused largely on other issues, such as the effect of real-world versus virtual reality testing (Waller et al., 2004), touching only briefly on construct validity. As will be discussed below, the JRD task is increasingly used in cognitive neuroscience experiments, in part due to its hypothesized connection to allocentric representation (Waller & Hodgson, 2006; Zhang et al., 2012, 2014; Ekstrom et al., 2014, 2017). Our findings here, first and foremost, support the overall construct validity of the JRD task, with some caveats. The results of our experiments support the use of the JRD task for studying spatial representations in cognitive neuroscience experiments. For example, we found that repetition enhances spatial memory precision and confidence. Previous cognitive neuroscience experiments have focused on studying changes in neural representations over the course of learning (Wolbers & Büchel, 2005; Suzuki & Brown, 2005; Komorowski, Manns, & Eichenbaum, 2009; Hargreaves, Mattfeld, Stark, & Suzuki, 2012; Deuker, Bellmund, Schröder, & Doeller, 2016). Thus, because performance changes over the course of the experiment, the combination of the navigation task and the JRD task can be used to study changes in spatial representations in brain areas known to be critical for spatial learning and memory, such as the hippocampus and retrosplenial cortex, as well as changes in interactions between such regions (Ekstrom et al., 2014, 2017).

Our results also suggest that participants are consciously aware of the precision of their spatial representations. These findings have potential application to real-world learning, including the enhancement of spatial knowledge. For example, it could be interesting to provide participants with increased exposure to the environment if they exhibit a low level of confidence overall. Taken further, it might be possible to provide participants with increased exposure to those landmarks for which they are least confident, or to provide them with feedback to help them improve their knowledge of the locations of such landmarks. Another interesting question, however, is whether such participants will have generally worse memory, regardless of overall exposure to the environment. In fact, some previous studies have suggested that participants with poor navigational ability, as assessed via questionnaire, tend not to improve on pointing tasks even after extensive exposure to the environment (Kozlowski & Bryant, 1977; Ishikawa & Montello, 2006). Moreover, in the present experiments, a notable proportion of participants failed to perform better than chance on the JRD task over the course of the experiment (Experiment 1: 8 of 24 participants; Experiment 2: 4 of 30 participants). These findings suggest that the JRD task is difficult, and future studies could address whether poorer performing participants could improve if they are provided with feedback. In fact, the findings from such manipulations may clarify how to improve spatial memory more generally, including in patients with memory-related disorders, such as Alzheimer’s disease (Viček & Laczó, 2014). In any case, the finding of individual differences in both performance and confidence will be important for studies investigating the neural involvement of different brain regions.

We found that the JRD task exhibits a high degree of test-retest reliability, thus supporting the construct validity of the JRD task. We also found a relationship between overall performance and test-retest reliability. These findings suggest that poorer performing participants might exhibit more variable spatial representations than better performing participants. As discussed above, this prediction could be tested by investigating whether there is a relationship between task performance and the variability of neural representations of spatial information, for example, by applying multivariate pattern analysis to fMRI or EEG data. Neuroimaging experiments typically require many trials and thus the JRD task is attractive because many trials can be presented. In fact, the JRD task has been used in previous fMRI experiments, revealing region-specific differences in blood-oxygen-level-dependent (BOLD) activity for the JRD task versus the SOP task (Zhang et al., 2012) and other tasks (Dhindsa et al., 2014), a relationship between hippocampal volume and performance on the JRD task (Schinazi et al., 2013), as well as differences in the brain regions coding for location and direction (Marchette, Vass, Ryan, & Epstein, 2014; Vass & Epstein, 2017). Thus, the JRD task has already proven useful in cognitive neuroscience experiments.

Taken together, our findings that repetition during navigation improves performance on the JRD task, that confidence ratings correlate with JRD performance, and that repeated trials on the JRD task result in stable answers over trials, suggesting its overall value as a stable and meaningful measure of spatial representations formed during navigation. Our results also suggest that the JRD task and the map drawing task recruit similar cognitive representations. In fact, both a group-level analysis and a within-subjects analysis of patterns of errors supported this hypothesis. These results further imply that the JRD task provides information about holistic, allocentric representations of large-scale space. Future experiments could examine the similarity of representations employed across different spatial memory tasks. For example, Marchette, Vass, Ryan, and Epstein (2015) reported similar representations between interior and exterior images of landmarks, implying a generalized code for these landmarks. Similar methods could be applied to study the representations employed across different cognitive tasks, thus allowing the inference of generalized neural representations of spatial information.

### 7.2 Future Experiments Can Use Similar Methodology

We implemented two techniques that we believe will be useful for future experiments. First, we described an analysis in which we investigated the pattern of errors across the JRD task and the map drawing task, and our results suggest that participants recruit similar cognitive representations to solve the two tasks. The analysis of error patterns can be useful for future studies that investigate the cognitive representations subserving performance across cognitive tasks, and it could also be used to compare cognitive representations, as assessed via error patterns in behavioral data, with neural representations, as assessed via error patterns in neural data (cf. Walther, Beck, & Fei-Fei, 2012). Second, we presented two group-level permutation tests, which provide straightforward methods for assessing chance performance, even when chance performance is not known *a priori* or when chance performance can differ across participants). Thus, the permutation tests described here will be useful for future studies using the JRD task, and they could potentially be used in other studies implementing group-level permutation tests, such as in the analysis of fMRI or EEG data (e.g., Chen et al., 2011; Liang et al., 2013; Stelzer et al., 2013; Etzel, 2015; Huffman & Stark, 2017).

### 7.3 Caveats Regarding Chance Performance on the JRD Task

Based on the results of our simulations, the guessing data, and the “null representation” experiment (in which participants performed the JRD task with no knowledge of the environment), we advocate for the use of permutation tests to determine chance performance on the JRD task. However, we wish to acknowledge that there are a number of caveats that need to be considered both with the scope of our findings and how the JRD task is conducted more generally. First, at the group level, both the guessing data and the “null representation” experiment suggested that chance pointing accuracy can be better than 90 degrees; however, pointing accuracy in these groups was 81.8 and 83.8 degrees, respectively, which are (numerically) fairly modest deviations from 90 degrees. One potential conclusion, then, is that observing pointing performance that is better than 90 degrees, at least under the condition in which JRD questions are sampled randomly, is a modest achievement and can sometimes be confounded with differences in pointing directions (i.e., a bias toward pointing in the forward-facing direction).

Second, there are cases in which chance performance will be 90 degrees. For example, if participants distribute their answers uniformly between 0 and 360 degrees, then the triangle rule does not apply (see the orange distribution in Figure 7). Thus, although we found evidence to suggest that this is not the case in both the guessing data and the null representation data (i.e., pointing accuracy was better than 90 degrees, suggesting a forward-facing bias), we cannot conclude that previous studies that have compared group-level performance to 90 degrees have had chance performance at levels other than 90 degrees (e.g., because participants in these studies might have uniformly distributed their responses). Nonetheless, we think that it will be important for future studies to consider the distribution of participants’ responses when making such comparisons—both at the single-subject level or at the group level.

Third, if JRD questions are sampled such that the answers are drawn from a uniform distribution between 0 and 360 degrees, then the triangle rule does not apply, and other studies have constrained JRD questions to come from a more uniform distribution (e.g., Shelton & McNamara, 2001; Mou et al., 2007; Shelton & Marchette, 2010). However, constraining the questions in this manner could potentially have other side effects, such as reducing the randomness of the questions themselves or the locations from which such questions are sampled (e.g., the locations of imagined position might not be random throughout the virtual environment; for related arguments about the importance of randomness in the context of stimuli in cognitive neuroscience experiments see: Westfall, Nichols, & Yarkoni, 2016). Additionally, including more trials—as allowed by either full sampling or a random sampling of questions—can be beneficial for measuring other aspects of pointing performance (cf. Marchette & Shelton, 2010) and under such conditions pointing direction can be used as a control regressor (e.g., Marchette, Yerramsetti, Burns, & Shelton, 2011) and comparisons to chance can be made using the permutation tests described here. Therefore, overall, we advocate for using permutation tests both to exclude chance-performing subjects and to compare group performance to chance, rather than using a fixed chance-level of 90 degrees, because these techniques provide a means of comparing to chance performance regardless of biases in pointing directions in (random) guessing and potentially allow for more robustness and flexibility in experimental design.

## Acknowledgements

Research supported by grants from NSF (BCS-1630296) and NIH (R01NS076856) awarded to Arne D. Ekstrom. Research also supported by a Ruth L. Kirschstein Postdoctoral Individual National Research Service Award from the National Institute of Mental Health of the National Institutes of Health (F32MH116577) awarded to Derek J. Huffman. The content is solely the responsibility of the authors and does not necessarily represent the official views of the National Institutes of Health. We thank Rebecca Mata, Michelle Occhipinti, and Nikhil Jaha for assistance with data collection. We thank members of the Human Spatial Cognition Lab, especially Michael Starrett and Jared Stokes, for helpful conversations about this project.

## Footnotes

1 Our focus here is on pointing tasks and map drawing tasks, thus the discussion of distance estimation tasks is limited.

2 We will refer to such tasks as “the SOP task” because this task could theoretically be solved using a mixture of egocentric and allocentric information (Ekstrom et al., 2014, 2017).

3 In our simulations we used mean angular error for comparison to previous studies.

4 We thank an anonymous reviewer for recommending a paired t-test approach.

